# Simplified and Rapid Workflow Enhances Throughput of *De Novo* Sequencing of COVID-19 Neutralizing Antibodies

**DOI:** 10.1101/2024.08.09.607349

**Authors:** Yueting Xiong, Jin Xiao, Wenbin Jiang, Jingyi Wang, Qingfang Bu, Xiaoqing Chen, Yangtao Wu, Rongshan Yu, Quan Yuan, Ningshao Xia

**Affiliations:** State Key Laboratory of Vaccines for Infectious Diseases, Xiang An Biomedicine Laboratory, School of Public Health, National Institute for Data Science in Health and Medicine, Xiamen University, Xiamen, Fujian 361102, China

## Abstract

During the current COVID-19 pandemic, precise antibody sequencing is crucial for the rapid development of broad-spectrum or pan-β-coronavirus neutralizing antibodies (NAbs) to prevent new variants of the coronavirus and other highly pathogenic β-coronaviruses. However, mass spectrometry-based *de novo* sequencing remains challenging due to its high cost, low throughput, and unexpected missing of certain ion information. We developed an innovative approach using solely bottom-up to provide a rapid, robust, and refined solution for the current *de novo* sequencing challenges. The methodology, referred to as SP-MEGD for Single-Pot and Multi-Enzymatic Gradient Digestion, capitalizes on a five-protease gradient digestion by sampling every two hours for a total of 6 hours in an integrative reactor. The SP-MEGD method could engender numerous missed cleavage events and produce various peptide products of diverse lengths with overlapping stretches of residues, enabling efficient database-free *de novo* sequencing. Antibody assembly for single or mixed COVID-19-NAbs with publicly available sequences and commercial antibodies with unknown sequences was efficiently deciphered using our previously proposed optimization method, Fusion assembler. Our innovative study represents the first successful simultaneous discrimination and achieving over 99% accurate sequence coverage of a mixture containing three humanized COVID-19-NAbs (S2P6LH, BD5514LH, and BD5840LH). Furthermore, we utilized SP-MEGD and Fusion to sequence commonly used anti-CD8 and anti-CD4, and successfully confirmed their impact on immune cell clearance through in vivo experiments. Overall, our developed workflow demonstrates promise in facilitating the discovery and development of vaccines and antibody therapeutics while providing deeper insights into monoclonal antibodies (mAbs).

## INTRODUCTION

More than two years since its emergence, the COVID-19 pandemic caused by severe acute respiratory syndrome coronavirus 2 (SARS-CoV-2) continues to spread. Neutralizing antibodies (NAbs) have emerged as pivotal players in preventing and treating COVID-19^1–3^. However, the persistent emergence of new variants has led to widespread evasion of NAbs, presenting significant challenges to SARS-CoV-2 NAb therapeutics^4^. Given their prophylactic and therapeutic efficacy, there is a high demand for the clinical development of NAb drugs that are resistant to future variants^5, 6^. The precise determination of the primary amino acid sequence of antibodies is a critical step in antibody drug discovery. The information of antibody sequence is crucial for comprehending the structural foundation of antigen-antibody binding, recognition, and interaction^7^. Additionally, sequence-based recombination expression or engineering modifications can be utilized to achieve constant region substitution of antibodies from different species, thereby altering the specificity of antibody species and allowing for the design of diverse antibodies.

However, obtaining the complete monoclonal antibodies (mAbs) sequence remains challenging. Traditional hybridoma technology^8–10^, which involves antibody screening and sequencing, typically takes three to six months to complete, requiring considerable time and cost. Researchers have made progress in isolating B cells directly from blood or bone marrow, extracting DNA or RNA, and creating sequencing libraries using second-generation sequencing technology independent of hybridoma technology, reducing the timeline to as short as one month^11–13^. Despite these advancements, the time-consuming nature of these methods and the need for complementary information from both genetic and protein levels remain challenges that must be addressed. Additionally, issues such as bacterial contamination or reduced viability during the antibody screening process can make obtaining sequence information impossible, severely limiting the flexibility of antibody screening. Furthermore, antibodies are secreted in bodily fluids and mucus without direct connection to their producing B cells. This raises questions about the quantitative relationship between the secreted antibody pool and the underlying B-cell population and potential sampling biases in current antibody sequencing strategies.

Mass spectrometry (MS)-based *de novo* sequencing of secreted antibodies serves as a valuable complementary means that can address several challenges confronted by conventional strategies relying on cloning/sequencing of the coding mRNAs^14, 15^. For example, the MS-based *de novo* sequencing approach can directly target polypeptide products in bodily fluids to acquire antibody sequences. It has successfully sequenced mAbs from lost hybridoma cell lines, establishing the foundation for the next generation of serological multi-antibody *de novo* sequencing. Nevertheless, despite advancements, MS-based *de novo* sequencing still encounters significant impediments such as large sample requirements, low throughput, unanticipated missing ion information, and difficulties in sequencing and assembly accuracy. During antibody sequencing, the existence of isomers with similar mass (e.g., leucine and isoleucine) or amino acid combinations (e.g., AG = Q, GG = N) can readily result in incorrect sequence assembly^16–18^. These problems are already encountered when sequencing a single antibody and become even more challenging for the *de novo* sequencing of multiple antibodies. Hence, choosing the correct peptides from ambiguous sequences and implementing precise amino acid error correction becomes crucial for developing more accurate, efficient, high-throughput antibody sequencing methods.

To overcome these limitations, we have innovatively proposed a fast, robust, and refined solution known as SP-MEGD (Single-Pot and Multi-Enzymatic Gradient Digestion). This approach employs a five-protease gradient digestion, with samples collected every two hours over a total duration of six hours within an integrated reactor. Subsequently, only the bottom-up tandem MS (MS/MS) method was utilized for data acquisition. This method has been shown to be applicable to antibodies with a sample volume as low as 50 μg. It leads to the production of multiple peptide products of diverse lengths, thereby increasing the overlapping extension of residues. By using the previously proposed optimization assembler Fusion in combination with SP-MEGD, it is feasible to efficiently and without reliance on a database, *de novo* sequence the entire heavy chain (HC) and light chain (LC) assembly of human COVID-19 neutralizing monoclonal antibodies (S2P6LH). Furthermore, we applied the SP-MEGD method to sequence commonly used anti-CD4 and anti-CD8 mouse monoclonal antibodies for which publicly available sequences are not accessible^19–21^. The technique achieves full sequence coverage of the HC and LC variable domains, including all complementarity determining regions (CDRs). The accuracy of the sequences obtained through experiments has been successfully verified through activity and function experiments. Additionally, our innovative study also encompassed identifying mixtures containing three human COVID-19 NAbs (S2P6LH, BD5514LH, and BD5840LH), achieving the simultaneous discrimination of three distinct antibodies for the first time with a sequence accuracy of over 99% each. This approach proves particularly beneficial in the context of pandemic infectious diseases as it enhances the efficiency and simplicity of identifying neutralizing antibody sequences, thereby accelerating the development of therapeutic and preventative measures. Furthermore, this approach holds potential in enabling a rapid response to newly emerging pathogens by generating targeted therapeutic antibodies or informing vaccine design strategies.

## EXPERIMENTAL SECTION

### Antibody Source

The anti-mouse CD8 (clone 2.43) and anti-mouse CD4 (clone GK1.5) monoclonal antibodies were purchased from Bio X Cell (BE0061 & BE0003-1). Our laboratory purified three humanized COVID-19 neutralizing antibodies (S2P6LH, BD5514LH, and BD5840LH) through the ExpiCHO-S™ suspension expression system (ThermoFisher, Cat#A29133). For further particulars, see the supporting information (SI).

### Sample Preparation Based on the SP-MEGD Method

Initially, the lysis buffer [6 M guanidine hydrochloride (Gu.HCl), 20 mM dithiothreitol (DTT), 100 mM Tris, pH 8.5] was added to 200 μg of purified monoclonal antibody (S2P6LH) at an approximate ratio of 1:1 (microliters of lysis buffer to micrograms of protein). The mixture was denatured and reduced at 60℃ for 30 min followed by alkylation with 40 mM iodoacetamide (IAA) in the dark for 30 min at room temperature. The alkylated antibody sample was transferred into a solution containing 0.8 M Urea, 50 mM Tris-HCl, pH 8.0 in a 10k filter (Millipore, USA) at 4 °C to eliminate interfering substances that might influence enzymatic digestion. The proteolytic enzymes, namely trypsin, chymotrypsin, pepsin, elastase, and Asp-N, were added at a ratio of 1:20 (w/w). The digestion solution was incubated at 37 °C for 6h, and samples were collected at two-hour intervals respectively. After digestion, the reaction mixture was quenched with 10% trichloroacetic acid (TFA) for 30 min at 37 °C. The supernatant was desalted by Sep-Pak C18 Vac cartridges (Waters) in accordance with the instructions before being lyophilized to dryness or stored at –20°C or dissolved in 0.1% FA for *de novo* LC−MS/MS analysis.

### *De Novo* LC–MS/MS Analysis

The digested peptides were separated through reversed-phase chromatography on a Vanquish™ Neo UHPLC (column packed with PepMap™ 100 C18, 75 μm×50 cm, 2 μm, Thermo Fisher Scientific, USA) coupled to an Orbitrap Eclipse mass spectrometer. Samples were eluted over a 70-min gradient from 0 to 35% solvent B (0.1% formic acid in 80% acetonitrile) at a flow rate of 300 nL/min. Solvent A was 0.1% formic acid in water. Full MS1 scans were acquired over a range of m/z 350−2000 with a resolution of 120, 000. MS1 scans were obtained with a standard automatic gain control (AGC) target and a maximum injection time of 100 ms. The precursors were fragmented by stepped high-energy collision dissociation (HCD) as well as electron-transfer high-energy collision dissociation (EThcD). The stepped HCD fragmentation included steps of 27%, 35%, and 40% normalized collision energies (NCE). EThcD fragmentation was performed with calibrated charge-dependent electron-transfer dissociation (ETD) parameters and 30% NCE supplemental activation. For both fragmentation types, MS2 scans were acquired at a 30,000 resolution, a 5e4, AGC target, and a 250 ms maximum injection time. Other method details are presented in the Supplementary Information.

### Machine Learning Model and Assembly Algorithm

We trained the Casanovo model (v3.2.0)^22^ on high-resolution MS/MS data from the human proteome using the HCD library from MassIVE, which consists of 2,154,269 spectra derived from 1,114,503 unique peptides^23^. The spectra were split into training and validation sets at a ratio of 98:2 while ensuring that the split data sets did not share any common peptides. The Casanovo model was then trained for 20 epochs using its pre-defined parameters. After training, we executed the Casanovo model to determine *de novo* peptide sequences from MS/MS spectra at a precursor tolerance of 10 ppm and fragment mass tolerance of 0.02 Da. Carbamidomethylation of cysteine (C + 57.02 Da) was set as a fixed modification, while oxidation of methionine (M + 15.99 Da) and deamidation of asparagine and glutamine (N + 0.98 Da and Q + 0.98 Da) were set as variable modifications. *De novo* peptides were then used by the Fusion assembler to automatically reconstruct the complete sequences of mAbs and mixtures of various mAbs. Human germline antibody sequences from IMGT are employed as templates to guide the assembly of *de novo* peptide reads.

### Intact Mass Verification of Antibody Light and Heavy Chain

One hundred micrograms of each antibody were reduced with 50 mM DTT and treated with 2 µl PNGase F (Promega, USA) at 37°C for over 16h with stirring before being transferred to a new vial for LC-MS analysis. More experimental conditions are given in the SI. **LC-MS Analysis of Intact Light and Heavy Chains.** Intact LCs and HCs were analyzed using an Orbitrap Eclipse mass spectrometer (Thermo Fisher Scientific, USA) coupled to a Vanquish™ Flex UHPLC (Thermo Fisher Scientific, USA). All details can be found in the SI.

### Data Analysis

For bottom-up analysis, the raw files obtained for digesting peptides of each antibody were searched with PEAKS AB (v. 3.0) using the parameters described in the SI. The intact mass spectra were processed and deconvoluted using BioPharma Finder 5.2 software (Thermo Fisher Scientific, USA), subsequently enabling the comparison of mass information for light and heavy chains.

### Expression and Purification of anti-CD8 and anti-CD4

The mAb sequences obtained from MS experiments and the Fusion method were utilized to generate antibodies in ExpiCHO-S cell following the manufacturer’s protocol. The proteins were purified from culture supernatants collected on day 7 after transfection using a Protein A column (Cytiva). Elution of the protein from the column was carried out with 70 mM citric acid monohydrate and 20 mM sodium phosphate dibasic dodecahydrate. The eluted fractions were pooled and concentrated using a Protein Quantification Kit (Thermo Fisher Cat#23225). SDS-PAGE analyses were performed using 4-12% SurePAGE precast mini polyacrylamide gels (Genscript), and gel images were captured using the FUSION FX7 Spectra multispectral imaging system (Vilber).

### Functional Validation of anti-CD8 and anti-CD4

Three groups of C57BL/6J mice (n=3 per group) were intraperitoneally injected with either 200 μg antibodies or 200 μL PBS buffer to observe the effective clearance rate of CD8^+^/CD4^+^ T cells in their PBMCs and spleens. The experiment was divided into three groups: PBS buffer, commercial anti-CD8/CD4 antibody, and anti-CD8/CD4 developed in our lab. Each group of mice was of the same age. On the first day, the mice received the antibody or PBS buffer injections. On the third day post-injection, single cells were isolated from peripheral blood or spleen tissue and resuspended in an appropriate volume of PBS buffer. Cell counting was performed, followed by treatment with a blocking buffer to reduce nonspecific antibody binding. Subsequently, the cell suspension was incubated with fluorescently labeled anti-CD8 and anti-CD4 antibodies. After incubation, the cells were washed with PBS buffer to remove unbound antibodies. The labeled cell suspension was then processed using a flow cytometer to measure the fluorescence intensity of the labeled dyes. Utilizing this data, CD8^+^ T cells and CD4^+^ T cells were sorted. Finally, FlowJo (BD Biosciences, V10.10) was used to analyze the data and assess the quantity, activity, and expression of the CD8^+^ and CD4^+^ T cells.

## RESULTS AND DISCUSSION

### Development and Optimization of the SP-MEGD Method

To establish a rapid, robust, and refined solution for single or mixed antibody *de novo* sequencing, we developed a methodology called SP-MEGD (Single-Pot and Multi-Enzymatic Gradient Digestion). This method leverages a five-protease gradient digestion approach, with samples taken every two hours over a total of 6 hours in an integrative reactor, aiming to minimize sample loss and provide abundant antibody ion information.

To develop the SP-MEGD, we rigorously optimized and evaluated several crucial aspects of the antibody preparation using the S2P6LH antibody with published sequences^24^. Our goal was to identify the optimal combination of reagents to streamline the protein denaturation and reduction steps, minimizing the sample loss in comparison to the conventional five-step process^25^. We compared the lysis efficiencies of three compatible lysis buffers: a. 2% DOC + 20 mM TCEP + 200 mM Tris-HCl, pH 8.0; b. 8 M Urea + 20 mM TCEP + 100mM Tris-HCl, pH 8.0; c. 6M Guanidine HCl + 20 mM DTT. The results were assessed based on protein sequence coverage (**Figure 2A**). It was observed that in HCD fragmentation mode, the coverage rates for heavy chains obtained from a combination of guanidine hydrochloride (Gu.HCl) buffer with various enzymes – trypsin, chymotrypsin, pepsin, elastase, and Asp-N – were 92.60%, 97.30%, 99.80%, 99.30%, and 96.20% respectively, all exceeding those achieved with other buffers. Similar trends were also seen in the EThcD fragmentation mode (**Figure 2A**). However, the coverage rates for the light chain from the combination of Gu.HCl buffer and Asp-N were not high, possibly due to fewer Asp-N cleavage sites in the tested antibody light chain sequence resulting in limited peptide information. Consequently, Gu.HCl buffer was subsequently selected for lysing antibody samples considering its overall effectiveness in achieving high coverage.

**Figure 1.**
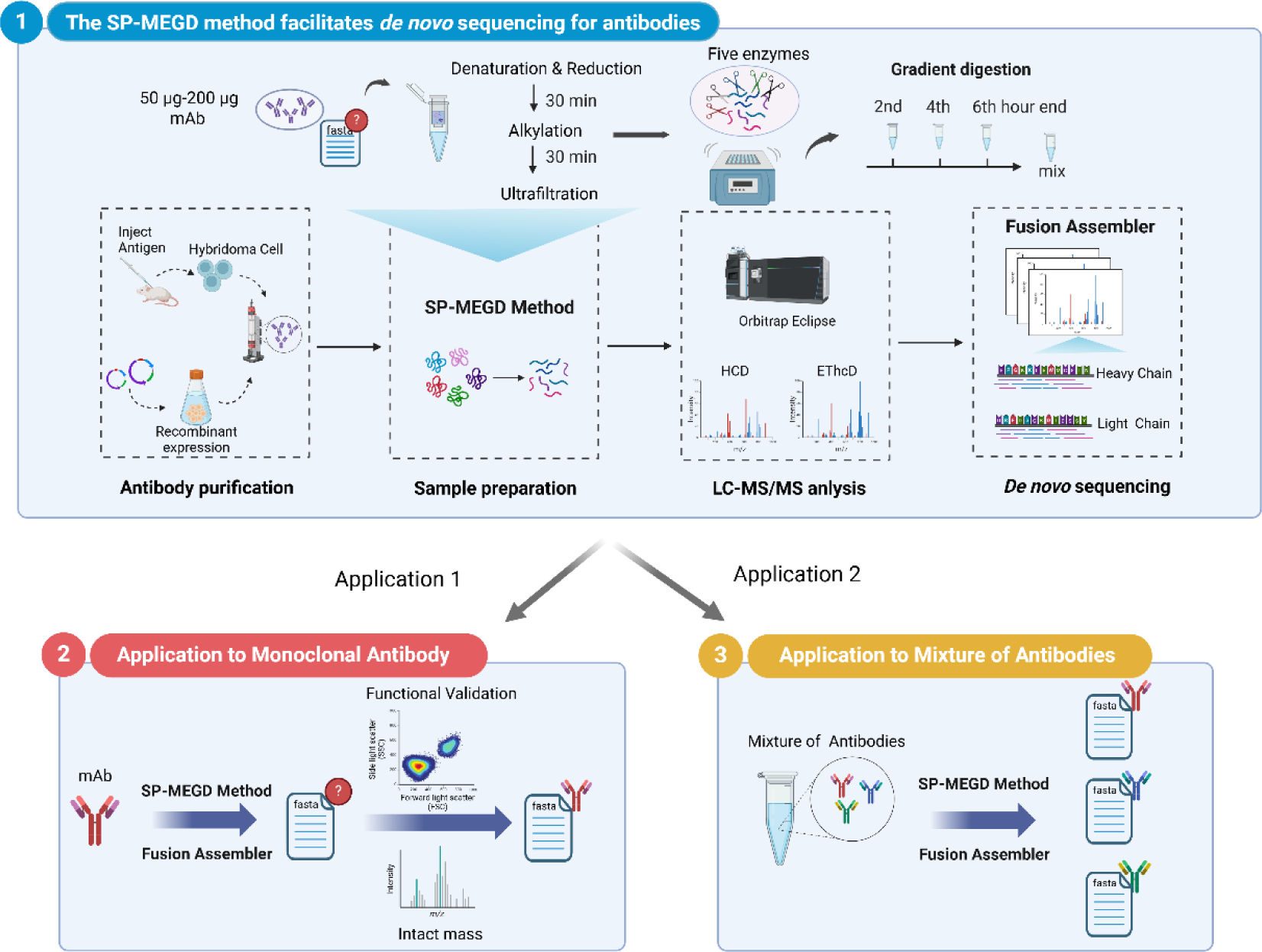
Framework for antibody *de novo* sequencing based on SP-MEGD preparation and Fusion assembler. Monoclonal antibody samples, approximately 50-200 µg each, were processed in a single pot with three main steps: denaturation and reduction for 30 minutes, followed by alkylation for 30 minutes, and then ultrafiltration. Subsequently, a gradient enzymatic digestion reaction was carried out using a combination of five enzymes. Samples were collected at the 2nd, 4th, and 6th hours before being pooled together. Purified peptides were identified using a dual fragmentation strategy (HCD & EThcD). Subsequently, *de novo* sequencing was performed using Casanovo model and Fusion assembler. Finally, the results were validated through intact mass analysis and bioactivity experiments.

**Figure 2.**
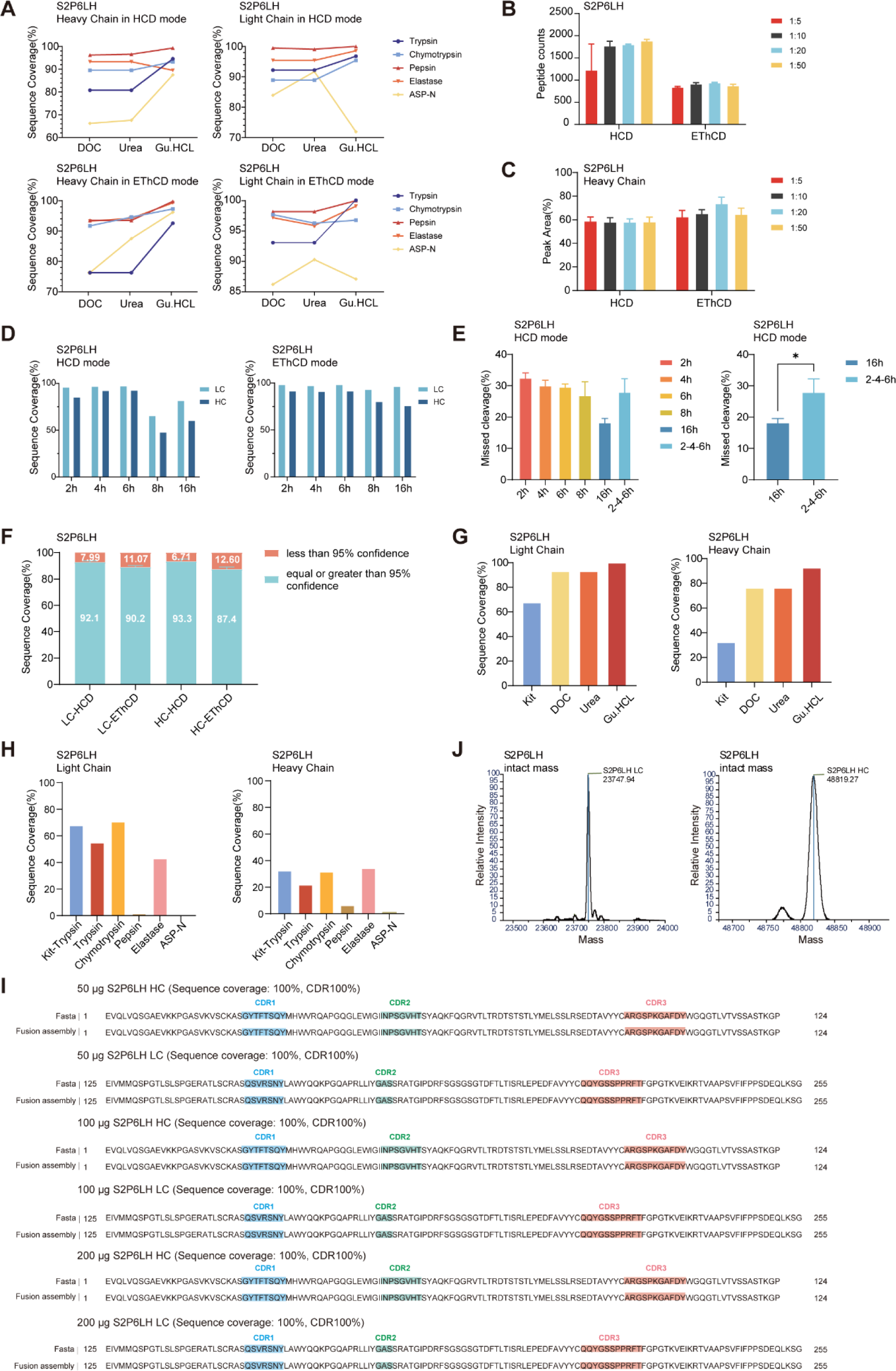
Development and Optimization of the SP-MEGD Method. (A) The sequence coverage of S2P6LH heavy and light chains after preprocessing with three different buffers: DOC, Urea, and Gu.HCl, along with corresponding enzymatic digestion combinations and MS fragmentation modes. (B-C) Peptide counts (B), peak area (C) of S2P6LH heavy chain coverage determined by various sample-to-trypsin digestion ratios, including 1:5, 1:10, 1:20, and 1:50. (D) Sequence coverage results of S2P6LH at different digestion times, including 2 h, 4 h, 6 h, 8 h, and overnight. (E) Comparison of missed cleavage rates at different digestion times using trypsin. The left panel shows missed cleavage rates at single time points; the right panel compares missed cleavage rates between gradient digestion (sampling at 2nd-4th-6th hours) and overnight digestion (sampling at 16 hours). (F) At different confidence levels, the sequence coverage of both the light and heavy chains of S2P6LH was assessed using various fragmentation modes. The analysis utilized a trypsin/protein ratio of 1:20 (w/w), with samples collected at the 2nd, 4th, and 6th hours. (G) SP-MEGD outperformed the commercial enrichment kit. Sequence coverage of S2P6LH heavy and light chains processed with EasyPept-Micro kit or with three different buffers under gradient digestion conditions using trypsin. (H) Comparison of sequence coverage of S2P6LH heavy and light chains between trypsin reagent from the EasyPept-Micro kit and various commercial digestion reagents. (I) Assembly result comparison of antibody samples at 50 µg, 100 µg, and 200 µg, using assembly coverage of S2P6LH heavy and light chains and accurate coverage of CDR regions as evaluation metrics. (J) The measured mass of S2P6LH heavy and light chains.

Traditional proteomics typically involves long reaction times and high enzyme-to-substrate ratios, resulting in a preference for low missed cleavage counts. However, sufficient peptide information is essential for *de novo* sequencing. Using the monoclonal antibody S2P6LH as a case study, we tested four different enzyme-to-protein ratios ranging from 1:5 to 1:50 of trypsin with a fixed 16-hour reaction time to ensure adequate reproducibility of the reaction. LC-MS/MS analysis was conducted to evaluate the unique proteolytic peptide count, peak area, and sequence coverage. As a result, there was an observed increase in the number of unique peptides as the enzyme-to-substrate ratio decreased from 1:5 to 1:50 (**Figure 2B**). In HCD mode, the peptide counts were 1881 at a ratio of 1:50 compared to 1880 at a ratio of 1:20. Meanwhile, in EThcD mode, the highest average peptide count was 939 at a ratio of 1:20. As indicated in **Figure 2C**, the peak area at a ratio of 1:20 demonstrated superior performance compared to other ratios. Moreover, the coverage of both light and heavy chains of the S2P6LH antibody was similar when the enzyme-to-substrate ratio was set at either 1:20 or 1:50. Thus, the optimal trypsin ratio was determined to be 1:20 in this context. This approach yields higher peptide counts and peak area, thereby satisfying the need of overlapping segments for *de novo* assembly. Similar optimizations were carried out for the remaining four predicted enzymes, each leading to their optimal enzymatic ratio (**Figure S1**).

Prolonged digestion may lead to a preference for low missed cleavage counts, but an increased missed cleavage rate can facilitate the generation of various peptide products with different lengths and overlapping residue extensions^26^. In this study, we investigated the optimal enzymatic digestion time for antibody samples to enhance peptide diversity and obtain sufficient peptide information. A comparative analysis of the digestion process was conducted, ranging from 2h to 16h (overnight) for antibody samples. The results indicated that the antibody sequence coverage increased from 2h to 6h of digestion, but decreased at 8h and overnight digestion (**Figure 2D**). Recognizing the significance of abundant fragment information in supporting peptide assembly in antibody *de novo* sequencing, we proposed a “gradient digestion” strategy. Samples were collected at the 2nd, 4th, and 6th hours before being pooled together to maximize missed cleavage rates (**Figure 2E**). This strategy could provide effective “head-to-tail” evidential information for assembling peptide fragments.

While a sequence coverage of over 87% (with a confidence level of at least 95%) has already been achieved using the 1:20 single trypsin digestion in gradient digestion (**Figure 2F**), notable enhancements in peptide counts can be acquired by exploiting multiple-enzyme combinations. Hence, the SP-MEGD method employs a five-protease (trypsin, chymotrypsin, pepsin, elastase, and Asp-N) gradient digestion approach, with samples collected every two hours over a total duration of 6 hours. This strategy offers more diversity in the length of proteolytic peptides, thereby augmenting the probability of overlapping fragments necessary for *de novo* assembly. Additionally, it provides enhanced flexibility and universalizes the digestion scheme regardless of the protein sequence and properties.

We carried out additional examinations to evaluate the attributes of the SP-MEGD method by employing diverse amounts (50 µg, 100 µg, and 200 µg) of starting materials for antibody samples. As depicted in **Figure 2I**, the S2P6LH antibody consistently achieved 100% accuracy in amino acid sequence within all CDR regions at the initial quantities of 50 µg, 100 µg, or 200 µg. This was determined by matching the protein sequence with the reference sequence of the test antibodies. These results demonstrate that our SP-MEGD method is suitable for analyzing antibody products with a sample volume as low as 50 µg, suggesting its considerable potential for routine usage.

### SP-MEGD Outperformed the Commercial Enrichment Kit

We compared the SP-MEGD method with the commercial EasyPept-Micro kit (Omicsolution, #OSFP0001) for microscale label-free sample preparation. To meet the microscale material requirement of 10-30 µg, we utilized 25 µg of S2P6LH antibody as an example for comparison. The SP-MEGD method achieved approximately 100% sequence coverage for the heavy chain of S2P6LH and 92.6% for the light chain when employing trypsin, surpassing those obtained from the EasyPept-Micro kit or other lysis methods (**Figure 2G-H**). As antibodies *de novo* sequencing often involves various proteases with complementary specificities, we also assessed the performance of different enzymes in the SP-MEGD method or in conjunction with a commercial EasyPept-Micro kit. Peptide information obtained using Pepsin and Asp-N in the EasyPept-Micro kit was notably lower than that obtained with the kit’s own trypsin reagent. This difference may be attributed to this particular kit reagent being incompatible with enzymes requiring acidic reaction conditions. These findings demonstrate that the innovative SP-MEGD strategy is more cost-effective and efficient than the commercial EasyPept-Micro kit.

### SP-MEGD method combined with Fusion method for *De Novo* Sequencing and Assembly

The LC-MS/MS method has been optimized to obtain high-quality data necessary for *de novo* sequencing. A dual fragmentation scheme, encompassing both HCD and EThcD on all peptide precursors, was chosen^27–29^. The stepped HCD fragmentation involves three collision energies (27%, 35%, and 40%) to cover multiple dissociation regimes, while the EThcD fragmentation is particularly effective for higher charged states and provides complementary c/z ions for maximum sequence coverage. These methods are mutually complementary.

Furthermore, the acquired data were analyzed using a trained Casanovo model to determine *de novo* peptide sequences. To assemble these peptides, we developed the Fusion assembler for template-assisted assembly of mAbs and mixtures of various mAbs. Fusion is a parameter-independent assembly method that only requires inputs regarding the species of the antibodies to be detected and their count. The k-mer parameter, which defines the number of overlapping amino acids needed to concatenate two peptides, is set by default to 7 in order to balance accuracy and coverage. Setting it too long results in insufficient peptide coverage, while setting it too short causes repetitiveness in target sequences.

By utilizing the Fusion assembler, we successfully achieved complete sequence coverage in both heavy and light chains across the variable and constant domains of the S2P6LH antibody **(Figure 2I**). Additionally, intact mass analysis was conducted to validate peptide sequences derived from the experiment. The experimental measurement of the mass of both light and heavy chains is consistent with the theoretically calculated value (**Figure 2J & Table 1**), falling within a reasonable margin of error. Overall, our innovative workflow yielded highly accurate sequences for the antibody with known sequences.

**Table 1.** Calculated and Measured Mass of Antibodies.

| **Name** | **From** | **$\Delta$ (C -M mass) (Da)** | **Measured mass (Da)** | **Calculated mass (Da)** | **DSB** | **K-loss** | **Pyro-Glu** |
| --- | --- | --- | --- | --- | --- | --- | --- |
| S2P6LH HC | Fusion | -2.58 | 48819.33 | 48953.00 | -8.08 | -128.17 | NA |
| S2P6LH LC | Fusion | 0.39 | 23747.94 | 23752.37 | -4.04 | NA | NA |
| BD5514LH HC | Fusion | -2.74 | 49323.06 | 49473.60 | -8.08 | -128.17 | -17.03 |
| BD5514LH LC | Fusion | 0.48 | 23613.46 | 23617.98 | -4.04 | NA | NA |
| BD5840LH HC | Fusion | 0.19 | 49173.67 | 49327.13 | -8.08 | -128.17 | -17.03 |
| BD5840LH LC | Fusion | 0.18 | 24342.64 | 24346.86 | -4.04 | NA | NA |
| CD4 HC 1 | PEAKS AB | -40.78 | 49238.11 | 49335.60 | -10.10 | -128.17 | NA |
| CD4 HC 2 | Fusion | 2.78 | 49238.11 | 49235.33 | -8.08 | -128.17 | NA |
| CD4 LC 1 | PEAKS AB | -0.08 | 23297.75 | 23301.71 | -4.04 | NA | NA |
| CD4 LC 2 | Fusion | 0.08 | 23297.75 | 23297.67 | -4.04 | NA | NA |
| CD8 HC 1 | PEAKS AB | -89.53 | 49286.86 | 49828.04 | -10.10 | -128.17 | NA |
| CD8 HC 2 | Fusion | 10.56 | 49286.86 | 49433.57 | -8.08 | -128.17 | NA |
| CD8 LC 1 | PEAKS AB | 0.58 | 23351.21 | 23355.83 | -4.04 | NA | NA |
| CD8 LC 2 | Fusion | -0.58 | 23351.21 | 23354.67 | -4.04 | NA | NA |

### Combining SP-MEGD method with Fusion algorithm to Sequencing Monoclonal Antibody

The global impact of SARS-CoV-2 has underscored the crucial role of T cells, specifically CD4+ and CD8+ T cells, in the immune response against the virus. Understanding the long-term immune memory generated after SARS-CoV-2 infection is essential for predicting reinfection risk and developing effective public health policies. By analyzing the antibody response targeting CD4+ and CD8+ T cells, we can assess the strength and durability of immune memory following infection or vaccination^19–21^. To address this need, we utilized the SP-MEGD and Fusion assembler to determine the amino acid sequence of anti-CD8 and anti-CD4 as test cases. Despite the widespread use of anti-CD4 and anti-CD8 to identify and deplete the CD4+ and CD8+ T cells, the only publicly available HC sequences are found in a patent for anti-CD8^30^.

### Anti-CD8

For the anti-CD8, a total of 112,541 MS2 scans were obtained with an average local confidence (ALC), out of which 55,442 MS2 scans showed more than 50% ALC. The median depth of overall coverage for the heavy chain of anti-CD8 was 201.73, while for the light chain, it was 293.80 (**Figure 3A**). There was better depth of coverage in the light chain than in the heavy chain. Additionally, we compared the performance of *de novo* sequencing for Fusion with PEAKS AB, a well-known antibody assembly software^31, 32^. The MS-based sequences of anti-CD8 derived from our previously developed Fusion method are shown alongside the sequence from PEAKS AB and a patent^30^ with highlighted CDRs in **Figure 3B**. And the full MS-based anti-CD8 sequences can be found in FASTA format in **Figure S2**.

**Figure 3.**
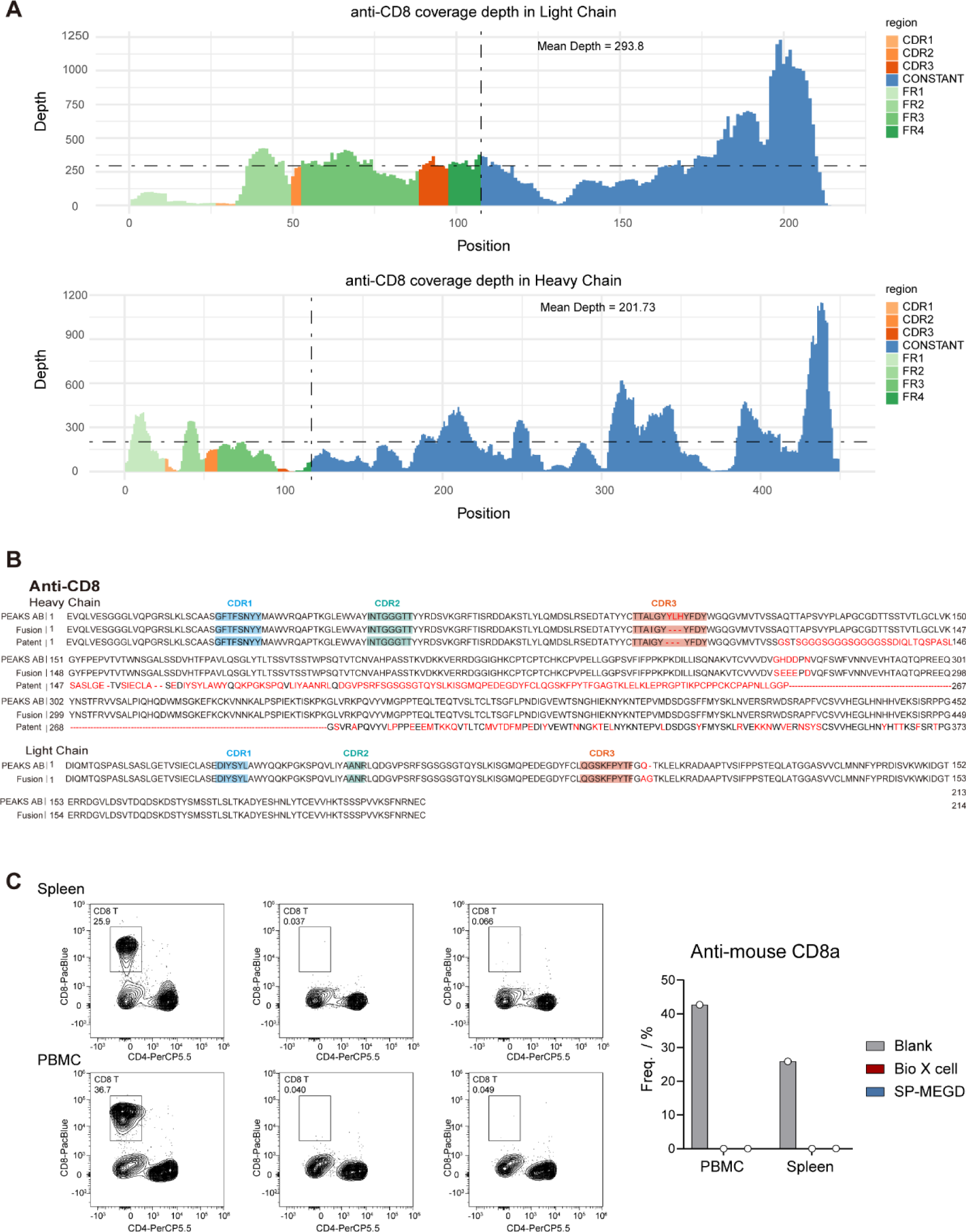
Combining SP-MEGD Method with Fusion algorithm to Sequencing Anti-CD8 Antibody. (A) A comparison of the coverage depth across different regions of the full-length light and heavy chains of the anti-CD8 antibody. The median total coverage depth for the anti-CD8 heavy chain is 201.73, while for the light chain, it is 293.80. (B) Alignment of the amino acid sequence of the anti-CD8 antibody using two assembly methods, PEAKS AB and Fusion. The heavy chain sequence is cross-referenced with published patent sequences. (C) A comparison of the efficiency in clearing CD8+ T cells in PBMCs and spleen across different anti-CD8 antibodies (Bio X cell and SP-MEGD), with a blank control group using PBS buffer.

We initially conducted the intact mass analysis to validate the experimentally determined sequences and compare the measured mass with the calculated mass derived from different sequences (**Table 1**). The mass discrepancy was smaller than 2 Da for light chains of anti-CD8, indicating an excellent match between the calculated and measured average masses. However, there was a Δmass larger than 2 Da for heavy chains of anti-CD8. Although the calculated mass of the light chain of anti-CD8 was consistent in both methods, we observed a different presence of amino acid combinations (AG = Q) in the two methods (**Figure 3B**). Additionally, there was a variation in the amino acid sequence in the CDR3 region of the heavy chain of anti-CD8 (**Figure 3B**). Therefore, further validation using biochemistry methods is warranted to confirm the accuracy of these two MS-based sequences.

Furthermore, the experimentally determined sequences from the Fusion method were cloned and transfected into cells for recombinant expression, followed by affinity purification. Subsequently, both the anti-CD8 developed in our lab (SP-MEGD) and a commercially available anti-CD8 (Bio X cell) were administered to C57BL/6J mice. Effective clearance of CD8+ T cells in both PBMCs and spleen was observed by the third day post-injection. When compared to control mice without T-cell depletion, the clearance efficiency of anti-CD8 (SP-MEGD) exceeded 95%, demonstrating comparable effectiveness to the commercial antibodies (**Figure 3C**). These findings suggest that our SP-MEGD method combined with the Fusion algorithm yields nearly complete antibody sequence coverage, providing high-confidence results for CDR regions across diverse mAb sequences. This could serve as a powerful tool for small-molecule drug screening in future applications.

### Anti-CD4

A total of 126,811 MS2 scans were identified for anti-CD4. The median depth of overall coverage of its heavy chain was 341.72, slightly higher than the light chain at 304.95 (**Figure 4A**). The MS-based sequences of anti-CD4 derived from the Fusion method were compared with the sequence from PEAKS AB, highlighting CDRs in **Figure 4B & Figure S3**. The light chain from both methods was found to be identical. However, the PEAKS AB method differed from the Fusion method in the experimentally determined sequence by 267 to 274 positions in the heavy chain of the constant region. Intact mass analysis was also conducted to validate the experimentally determined sequences derived from both methods. The difference in the light chain mass of anti-CD4 antibodies was only 0.08 Da between the calculated and measured mass, confirming the accuracy of the experimentally determined light chain. However, it is worth noting that the anti-CD4 heavy chain generated by the PEAKS AB method exhibited a Δmass of 40.78 Da, compared to 2.78 Da by the Fusion method. It is speculated that the sequence generated by Fusion may more faithfully represent the commercial anti-CD4 antibody. The efficacy of the anti-CD4 sequences obtained by SP-MEGD and Fusion methods was verified using an in vivo immune cell clearance assay (**Figure 4C**).

**Figure 4.**
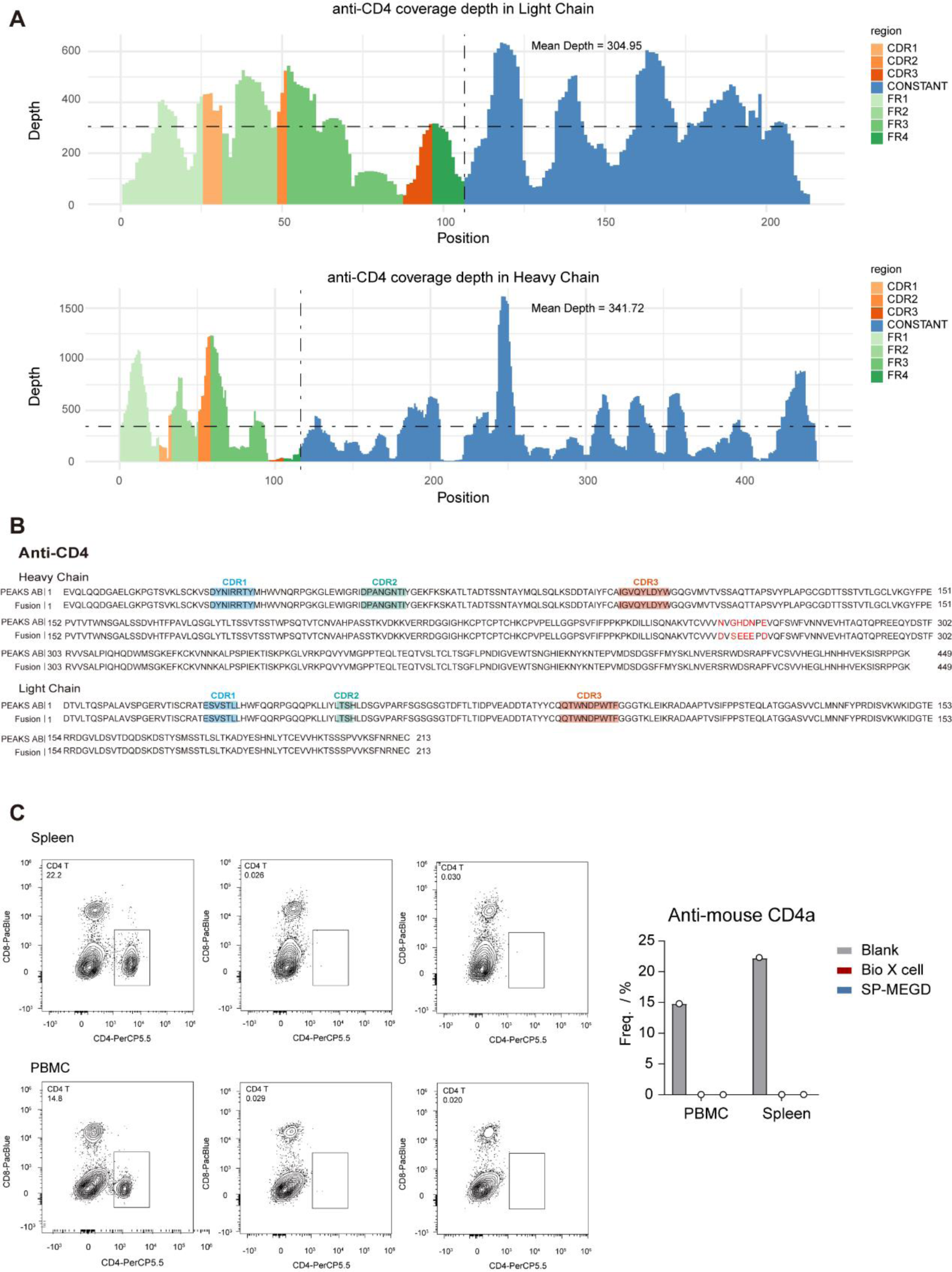
Application of SP-MEGD with Fusion Method to Anti-CD4 Antibody. (A) A comparison of the coverage depth across different regions of the full-length light and heavy chains of the anti-CD4 antibody. The median total coverage depth for the anti-CD4 heavy chain is 341.72, while for the light chain, it is 304.95. (B) Alignment of the amino acid sequence of the anti-CD4 antibody using two assembly methods, PEAKS AB and Fusion. (C) Validation of the biological activity of the anti-CD4 antibody.

### Combining SP-MEGD method with Fusion assembler to Sequencing the Mixture of COVID-19 Neutralizing Antibodies

In-depth research on neutralizing antibodies against COVID-19 not only provides a scientific foundation for current vaccine and therapeutic strategies, but also offers crucial scientific evidence for future efforts against the virus and its variants. Understanding and isolating effective neutralizing antibodies from convalescent patients can provide passive immune support to individuals in the early stages of the disease. Specific neutralizing antibodies have been utilized to develop monoclonal antibody drugs, demonstrating clinical efficacy in certain COVID-19 patients. Therefore, obtaining the primary sequences of these antibodies is essential. However, directly sequencing polyclonal antibodies derived from host serum poses significant analytical challenges, and analyzing a monoclonal antibody sequence can be time-consuming and costly. To address this issue, we aimed to test the applicability of our self-developed SP-MEGD method and Fusion assembler for sequencing mixtures of multiple monoclonal antibodies.

First, we mixed equal proportions of two antibodies, S2P6LH and BD5514LH. We conducted sample preparation using the SP-MEGD method and performed sequence assembly using the Fusion algorithm. The results demonstrated that the sequence accuracy for S2P6LH reached 100%, with a CDR region accuracy of 100% in both the heavy and light chains, as compared to the known original sequences (**Figure 5A**). For BD5514LH, the whole sequence accuracy of the light chain was 100%, while 99.56% of the heavy chain showed accurate sequencing (**Figure 5A**).

**Figure 5.**
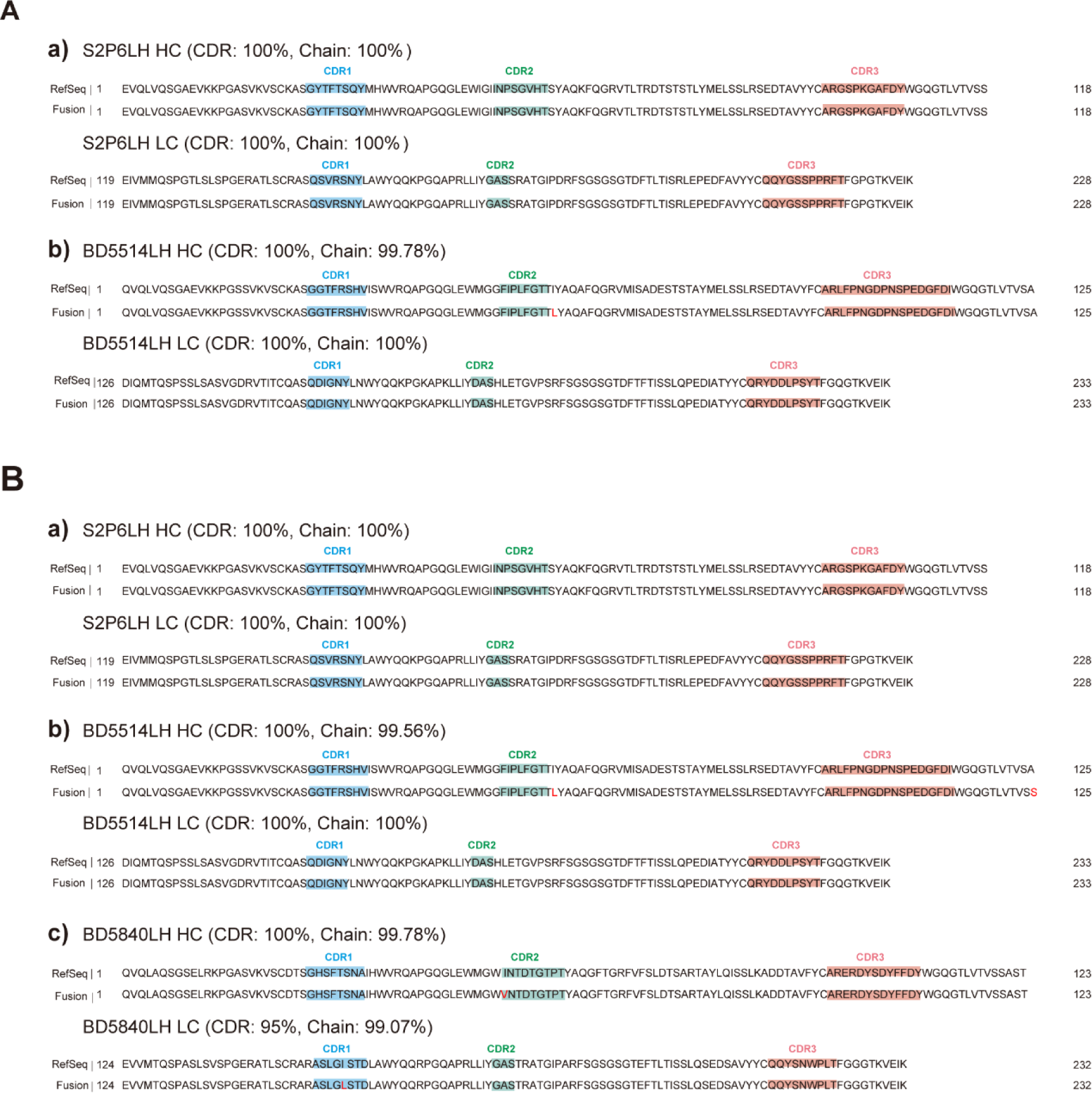
Application of SP-MEGD with Fusion Assembler to the Mixture of COVID-19 Neutralizing Antibodies. (A) Combining the SP-MEGD method with the Fusion algorithm, the experiment-determined light and heavy chain sequences of an equal mixture of two monoclonal antibodies, S2P6LH and BD5514LH, were compared to the known sequences. This comparison includes coverage of the full-length sequences and accuracy of coverage in the CDR regions. (B) Combining the SP-MEGD method with the Fusion algorithm, the experiment-determined light and heavy chain sequences of an equal mixture of three monoclonal antibodies, S2P6LH, BD5514LH, and BD5840LH, were compared to the known sequences. This comparison includes coverage of the full-length sequences and accuracy of coverage in the CDR regions.

Next, we mixed three antibodies in equal proportions. For S2P6LH, both heavy and light chain coverage reached 100%, with an accuracy of 100% in their respective CDR regions (**Figure 5B**). As for BD5514LH, the light chain achieved full coverage at 100%, with a CDR sequence accuracy also reaching 100%. However, there was a slight decrease in accuracy for the whole heavy chain at 99.78% (**Figure 5B**). This discrepancy may be attributed to increased difficulty in distinguishing between amino acids I and L when all three antibodies are mixed together. In regards to BD5840LH, both heavy and light chain coverage exceeded 99%, with CDR sequence accuracies of 100% for the heavy chain and 95% for the light chain. Notably, a difference was observed in the CDR2 region of the heavy chain for BD5840LH. The original sequence indicated an isoleucine (I), while all *de novo* sequences showed a valine (V). Suspecting a potential plasmid mutation, we sequenced the CDR region of the BD5840LH heavy chain. The sequencing results confirmed that the first 3’ to 5’ base pair in the CDR2 region was GAC, corresponding to valine (Val/V), making the sequence VNTDTGTPT. The provided theoretical base sequence was GAT, corresponding to isoleucine (Ile/I), with the sequence INTDTGTPT. Thus, the actual accuracy of the CDR sequence for the heavy chain of BD5840LH antibody was found to be 100%. This finding suggests that our developed method and algorithm can accurately assemble correct and authentic sequences effectively correcting amino acid mutations in antibody sequences. It substantiates the accuracy and effectiveness of our method and highlights its considerable potential for future applications.

## CONCLUSIONS

We have developed a simplified and rapid *de novo* sequencing workflow for the complete characterization of single or mixed COVID-19 neutralizing antibodies using publicly available sequences, as well as for commercial antibodies with unknown sequences. This workflow incorporates a Single-Pot and Multi-Enzymatic Gradient Digestion (SP-MEGD) method to provide abundant peptide information, and a beam search based assembler (Fusion) for antibody assembly.

By utilizing this pipeline, we were able to effectively characterize commercial antibodies (anti-CD8 and anti-CD4) for which with no publicly available sequences existed. The experimentally determined sequences were cloned and transfected into cells for recombinant expression, followed by affinity purification and functional validation. This demonstrated comparable effectiveness to the commercial antibodies. The availability of the anti-CD8 and anti-CD4 sequence may significantly contribute to the widespread use of this crucial research tool. This example serves as an illustration that our MS-based sequencing protocol produces robust and reliable antibody sequences. Furthermore, our MS-based SP-MEGD method and Fusion assembler were also applied in the sequencing of a mixture containing two or three COVID-19 neutralizing antibodies. The high-quality spectra obtained from the SP-MEGD method are suitable for *de novo* sequencing and may further advance the exciting prospect of a new advanced serology, in which antibody sequences can be directly obtained from bodily fluids.

## AUTHOR INFORMATION

### Corresponding Authors

**Yueting Xiong −** State Key Laboratory of Vaccines for Infectious Diseases, Xiang An Biomedicine Laboratory, School of Public Health, National Institute for Data Science in Health and Medicine, Xiamen University, Xiamen, Fujian 361102, China

**Rongshan Yu −** State Key Laboratory of Vaccines for Infectious Diseases, Xiang An Biomedicine Laboratory, School of Public Health, National Institute for Data Science in Health and Medicine, Xiamen University, Xiamen, Fujian 361102, China

**Quan Yuan −** State Key Laboratory of Vaccines for Infectious Diseases, Xiang An Biomedicine Laboratory, School of Public Health, National Institute for Data Science in Health and Medicine, Xiamen University, Xiamen, Fujian 361102, China

**Ningshao Xia −** State Key Laboratory of Vaccines for Infectious Diseases, Xiang An Biomedicine Laboratory, School of Public Health, National Institute for Data Science in Health and Medicine, Xiamen University, Xiamen, Fujian 361102, China

### Authors

**Jin Xiao −** State Key Laboratory of Vaccines for Infectious Diseases, Xiang An Biomedicine Laboratory, School of Public Health, National Institute for Data Science in Health and Medicine, Xiamen University, Xiamen, Fujian 361102, China

**Wenbin Jiang −** State Key Laboratory of Vaccines for Infectious Diseases, Xiang An Biomedicine Laboratory, School of Public Health, National Institute for Data Science in Health and Medicine, Xiamen University, Xiamen, Fujian 361102, China

**Jingyi Wang −** State Key Laboratory of Vaccines for Infectious Diseases, Xiang An Biomedicine Laboratory, School of Public Health, National Institute for Data Science in Health and Medicine, Xiamen University, Xiamen, Fujian 361102, China

**Qingfang Bu −** State Key Laboratory of Vaccines for Infectious Diseases, Xiang An Biomedicine Laboratory, School of Public Health, National Institute for Data Science in Health and Medicine, Xiamen University, Xiamen, Fujian 361102, China

**Xiaoqing Chen −** State Key Laboratory of Vaccines for Infectious Diseases, Xiang An Biomedicine Laboratory, School of Public Health, National Institute for Data Science in Health and Medicine, Xiamen University, Xiamen, Fujian 361102, China

**Yangtao Wu −** State Key Laboratory of Vaccines for Infectious Diseases, Xiang An Biomedicine Laboratory, School of Public Health, National Institute for Data Science in Health and Medicine, Xiamen University, Xiamen, Fujian 361102, China

### Author contributions

Y.X., J.X., and W.J. contributed equally to this work. This manuscript was written through the contributions of all authors. All authors have approved the final version of the manuscript.

### Notes

The authors declare no conflict of interest.

## ACKNOWLEDGMENTS

This work is supported by the Natural Science Foundation of Xiamen, China (3502Z202371039).

